# Metapredict V2: An update to metapredict, a fast, accurate, and easy-to-use predictor of consensus disorder and structure

**DOI:** 10.1101/2022.06.06.494887

**Authors:** Ryan J. Emenecker, Daniel Griffith, Alex S. Holehouse

## Abstract

Intrinsically disordered proteins and protein regions make up 20-40% of most eukaryotic proteomes and play essential roles in a wide gamut of cellular processes, from intracellular trafficking to epigenetic silencing. Given their importance, the ability to robustly, quickly, and easily identify IDRs within large proteins is critical. Here we present metapredict V2, an update to our deep-learning-based disorder predictor metapredict. Metapredict V2 has substantially improved accuracy, more features, and a more user-friendly interface via our web server (https://metapredict.net/), Python package, and command-line tool. To illustrate V2’s improved performance we undertake a systematic analysis of human transcription factors, as well as illustrate that metapredict V2 works well for synthetic or non-natural proteins.

**KEY POINTS:**

- Metapredict is a fast and easy-to-use disorder predictor released in 2021.
- Metapredict V2 was released in March 2022 and includes improved accuracy and new features.
- Metapredict V2 is now the default metapredict implementation, although the original implementation is available as ‘legacy’ metapredict.
- This manuscript provides a summary of how we improved the accuracy of metapredict and compares the original version (legacy) to our improved version (V2)
- This manuscript will not be submitted to a journal; if you use metapredict V2 please cite the original paper and make reference to the fact that V2 is being used.

## INTRODUCTION

In the summer of 2021 we published metapredict, a fast, accurate, and easy-to-use software tool for predicting intrinsically disordered regions (1). Over the last eight months, we have continued to improve metapredict’s features and accuracy, culminating in version 2 (V2) released in March of 2022.

This short article reports some of those improvements and explains several features that have changed since the original metapredict implementation (which we refer to as ‘legacy’ metapredict). These changes include improvements in accuracy, an improved web server, and a more intuitive interface for working with disordered domains. In addition, we performed a systematic analysis of all human transcription factors to excise their predictor IDRs and reveal the extent of predicted disorder within these proteins. We chose transcription factors because their amino acid sequence characteristics often raise challenges in robustly identifying IDRs. As such, we hope this new analysis will be useful for others working in the transcription field.

This article will exist as a preprint on bioRxiv, and we have no plans to submit it to a journal. If you find metapredict useful, please consider citing the original metapredict paper but clarify which version of metapredict you are using (V2 vs. legacy) in the methods section. We generally recommend V2; based on our updated benchmarking V2 offers a substantial improvement in the accurate identification of IDRs and better resolution of boundaries between IDRs and folded domains. The V2 predictor is now the default predictor, provided you are using metapredict version 2.0 or higher, both locally and with the metapredict web server. As reported in our original paper, legacy metapredict may be more conservative. However, we note one important caveat - if correlations with AlphaFold2 derived predictions are of interest, we recommend using legacy given that V2 integrates information from AlphaFold2.

## RESULTS

### Combining predicted pLDDT scores with metapredict disorder scores improves the accuracy of disorder predictions

The original release of metapredict included the ability to predict AlphaFold2 (AF2) per-residue predicted <u>L</u>ocal <u>D</u>istance <u>D</u>ifference <u>T</u>est (pLDDT) scores. In the context of AF2, pLDDT scores convey the confidence associated with a given structure prediction (2). Using the original AF2 data, we trained a bidirectional recurrent neural network with long short-term memory (BRNN-LSTM) using the generalized deep learning package PARROT. This enabled the prediction of pLDDT scores from sequence alone (3). Although low pLDDT scores cannot strictly be interpreted directly as a given region having a high likelihood of being disordered, low pLDDT scores generally correlate with disorder scores (1, 2, 4, 5). Therefore, our original release’s inclusion of predicted pLDDT (ppLDDT) scores provided an orthogonal means to examine whether a protein region is likely to be disordered.

Shortly after the release of metapredict, we began examining ways of combining our metapredict predicted consensus disorder scores with ppLDDT scores such that we could improve the accuracy of predicted disordered regions generated by metapredict. The most successful approach involved combining inverse ppLDDT scores normalized to values between 0 and 1 with metapredict consensus disorder scores. In this implementation, if either value was above 0.5, that value was used. If both the metapredict consensus disorder score and the normalized inverse ppLDDT score were below 0.5, the lower value between the two was used. This predictor, which we call ‘metapredict hybrid,’ resulted in a substantial increase in accuracy compared to our original implementation (‘legacy metapredict’) (**Fig. 1A**). Moreover, the resulting disorder profiles were much easier to interpret than those generated by legacy metapredict, with ordered and disordered domains clearly delineated (**Fig. 1B, C**). These results confirmed that incorporating ppLDDT information into metapredict improves accuracy and interpretability.

**Figure 1.**
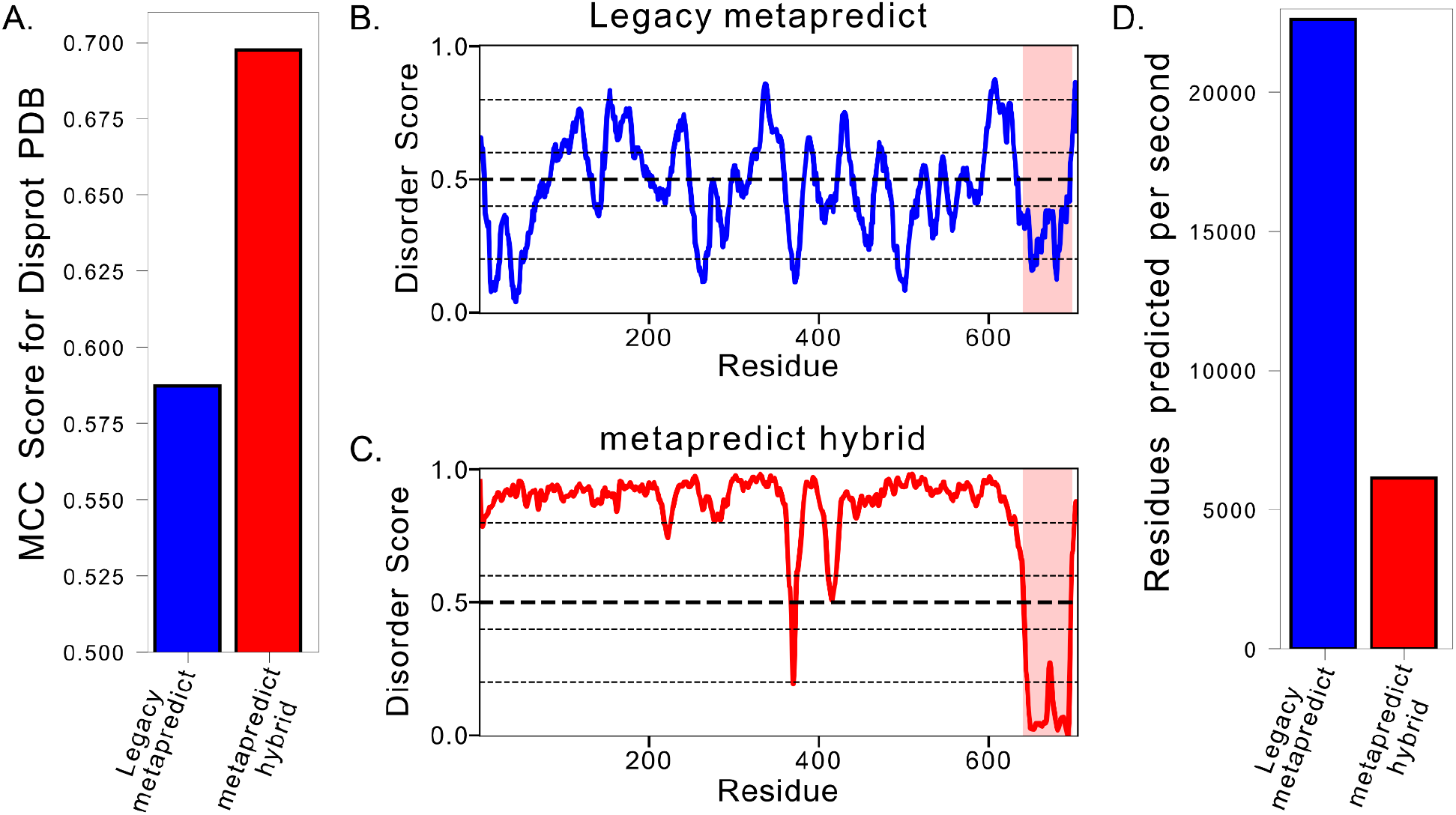
Comparing the original metapredict disorder predictor to metapredict hybrid. **(A)** Mathew’s correlation coefficient (MCC) scores for binary disorder predictions using the Disprot-PDB dataset (6). The **(B)** Predicted disorder for *Saccharomyces cerevisiae* Msn2 (Uniprot ID - P33748) using the original implementation of metapredict. The Zn-finger-containing DNA binding domain (DBD) is highlighted in red. **(C)** Predicted disorder for the same protein as in (B) using the metapredict hybrid disorder predictor, with the DBD again highlighted. **(D)** Assessment of the number of residues predicted per second using 500 randomly generated sequences of 500 amino acids in length.

Despite these improvements, metapredict hybrid was substantially slower than legacy metapredict (**Fig. 1D**). While legacy metapredict uses a single-layer BRNN-LSTM, metapredict hybrid makes sequential calls to two separately trained single-layer networks and then combines the results. Metapredict hybrid, therefore, offered improved accuracy at the expense of slower execution times.

### Training a new neural network on metapredict hybrid disorder scores to increase disorder prediction speed without losses in accuracy

Given that an important feature in the legacy metapredict implementation was its execution speed, we sought to develop a method to retain the improved accuracy of metapredict hybrid while minimizing the slower execution speeds. To this end, we generated metapredict hybrid scores for 363,265 protein sequences from 21 proteomes, which we then used to train a new BRNN-LSTM that could recapitulate metapredict hybrid scores (**Fig. 2**). Training, testing, and validation were done using our deep learning toolkit PARROT (3). In training this new network, we tested several distinct combinations of hyperparameters and found that a BRNN-LSTM with two hidden layers was necessary to recapitulate the original training data accurately. This was in contrast to legacy metapredict, which only needed a single layer.

**Figure 2.**
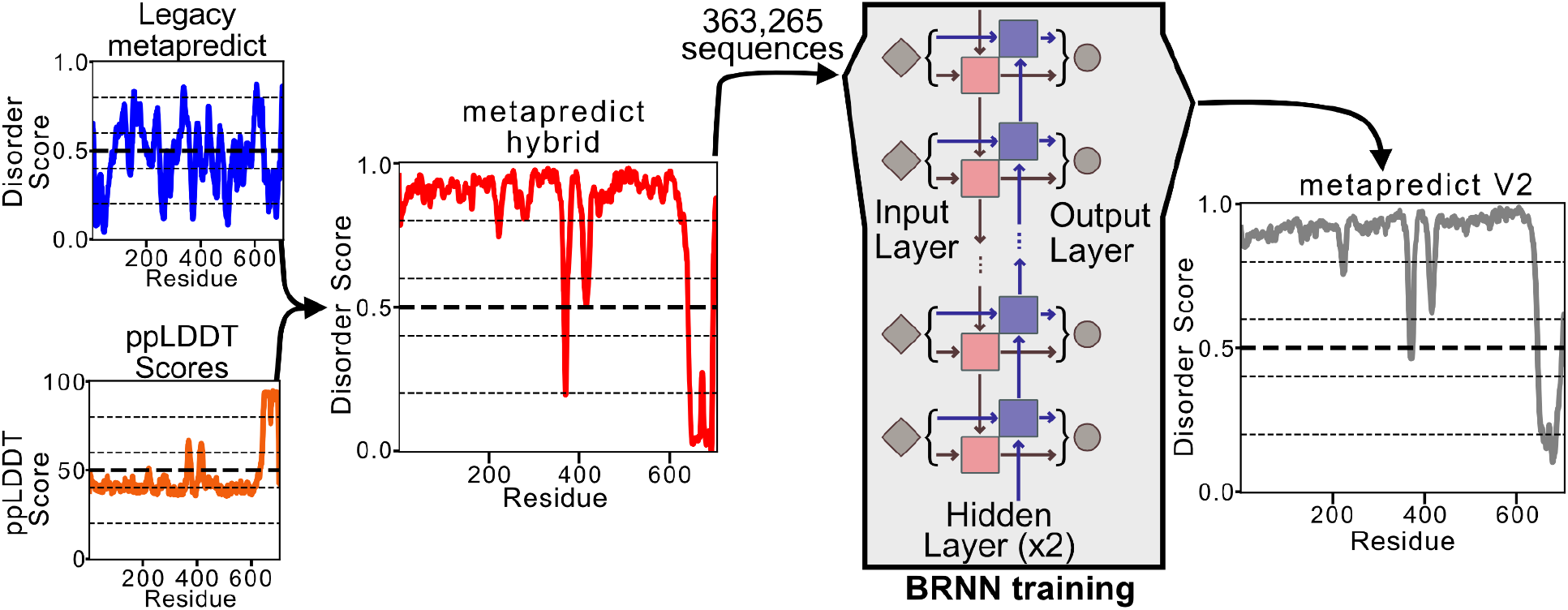
Workflow for the creation of metapredict V2. Predicted pLDDT (ppLDDT) scores were combined with disorder scores from the original version of metapredict (legacy metapredict) to generate metapredict hybrid scores. Metapredict hybrid scores were generated for 363,265 sequences, which were used to generate a bidirectional recurrent neural network (BRNN) that is used to generate disorder scores for metapredict V2.

When we assessed the accuracy of this newly trained predictor (metapredict V2) using the Disprot-PDB dataset (as done previously (1, 7, 8)), we found that it was slightly more accurate than metapredict hybrid and much more accurate than legacy metapredict (**Fig. 3A**) (1). Importantly, although metapredict V2 is still slower than legacy metapredict, it is substantially faster than metapredict hybrid (**Fig. 3B**). We attribute this difference in performance to the difference in hidden layers between legacy and V2 (1 vs. 2). Finally, when we analyzed disorder across individual proteins, the disorder profiles generated by metapredict V2 and metapredict hybrid were nearly identical (**Fig. 3C**). In summary, metapredict V2 offers all the benefits of hybrid-metapredict with a slight increase in accuracy and a substantial increase in execution speed.

**Figure 3.**
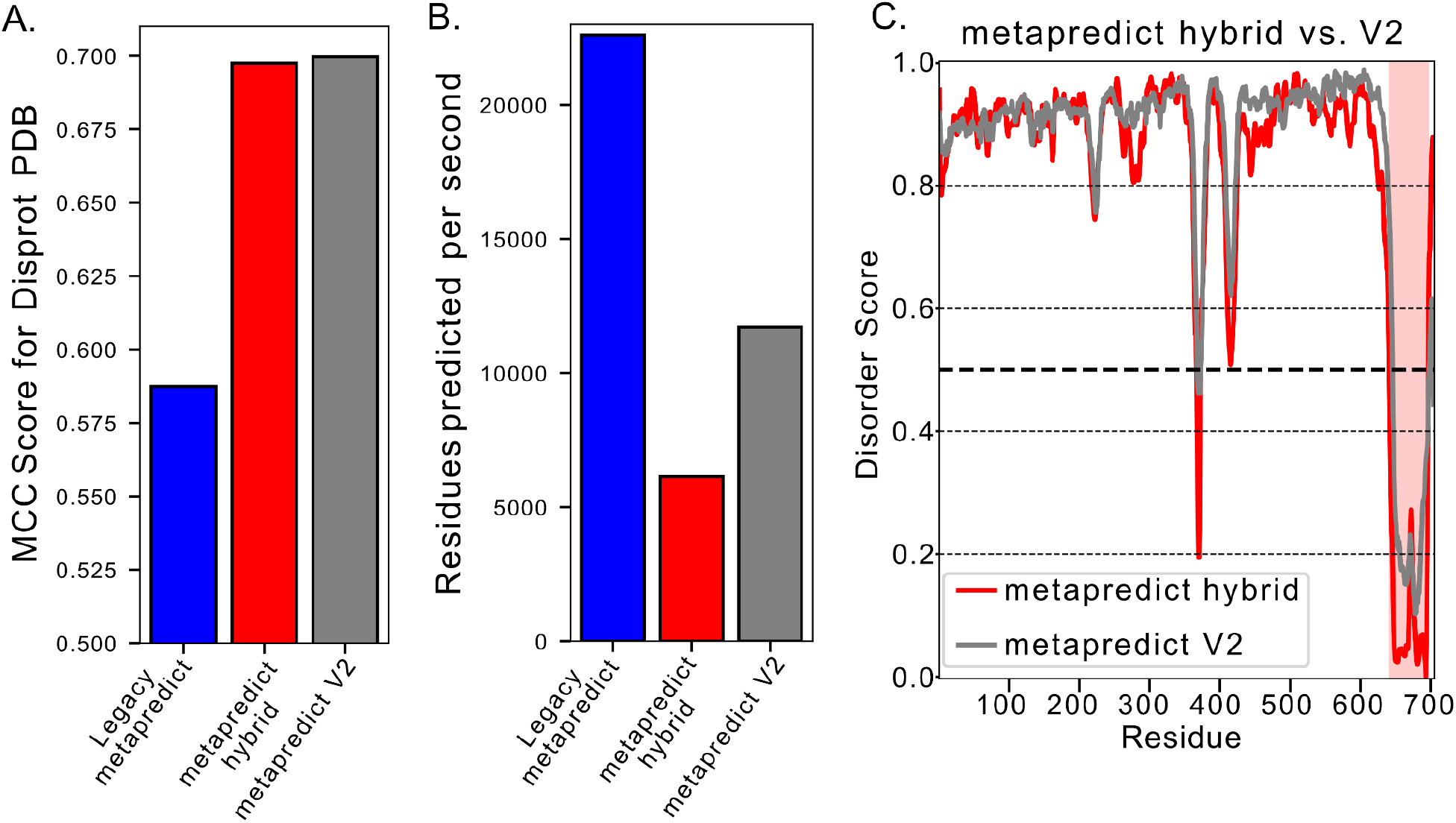
Metapredict V2 has comparable accuracy to metapredict hybrid with faster execution times. **(A)** Mathew’s correlation coefficient scores (MCC) for binary disorder predictions using the Disprot-PDB dataset. A threshold of 0.5 was used to define if a residue was ordered or disordered. **(B)** Assessment of the number of residues predicted per second using 500 randomly generated sequences of 500 amino acids in length. **(C)** Predicted disorder for *Saccharomyces cerevisiae* Msn2 (Uniprot ID - P33748) comparing metapredict hybrid to metapredict v2. The Zn-finger-containing DNA binding domain (DBD) is highlighted in red.

### Metapredict V2 more accurately identifies fully disordered proteins than legacy metapredict

Although legacy metapredict was relatively accurate and extremely fast, we were able to find examples where metapredict was unable to identify disordered regions that had previously been experimentally characterized as disordered. Some of these examples were identified in publications that used metapredict (4), whereas others were identified by examining fully disordered proteins that legacy metapredict failed to identify as disordered within the Disprot-PDB dataset. In examining the predicted disorder for five of these proteins, we found that metapredict V2 did substantially better at identifying these proteins as disordered than did legacy metapredict (**Fig. 4A, B**).

**Figure 4.**
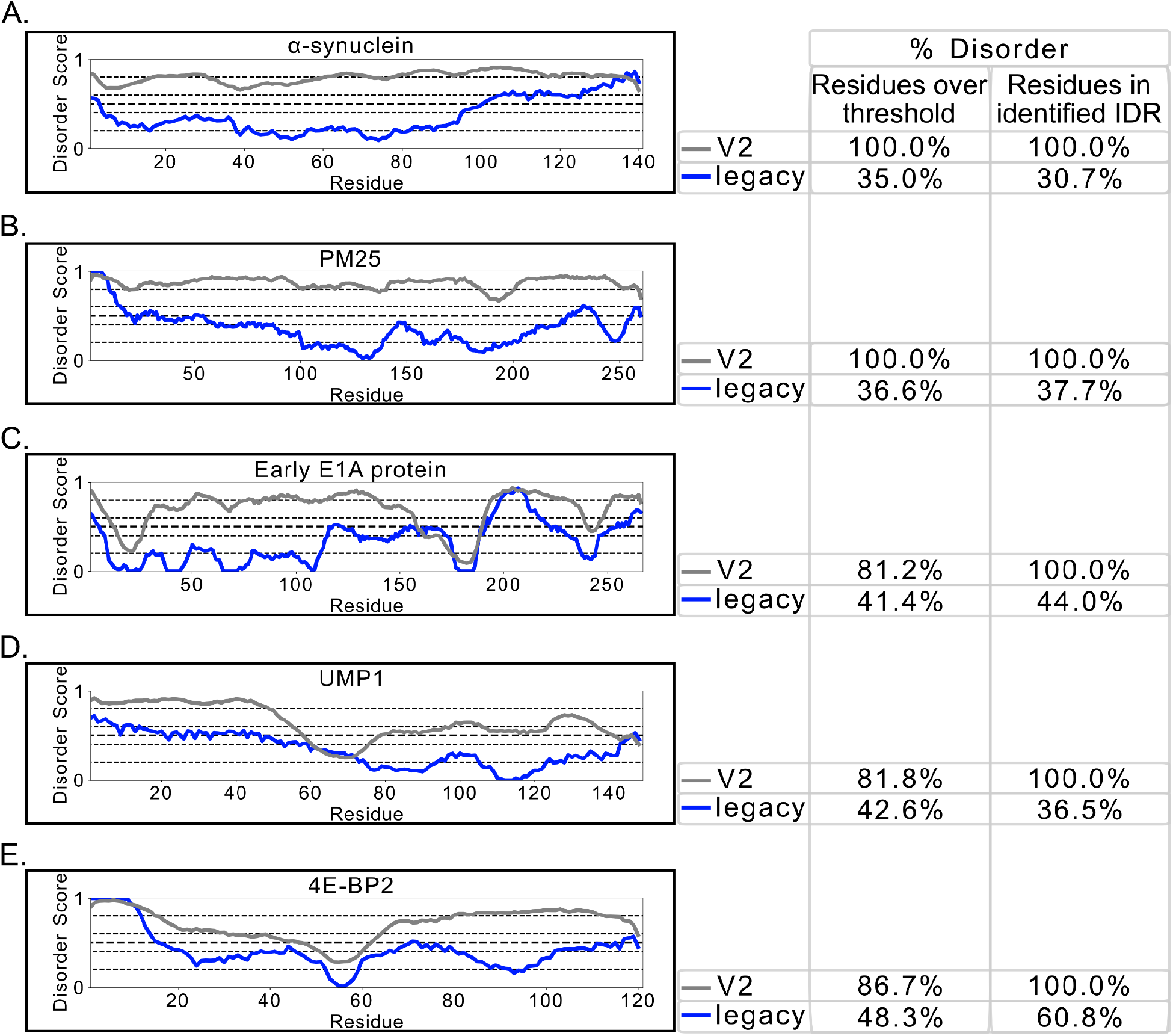
Metapredict V2 more accurately identifies disorder in experimentally determined disordered proteins or disordered protein regions in comparison to legacy metapredict. Each graph (left) shows the disorder profiles generated using either metapredict V2 or legacy metapredict for the proteins **(A)** *H. sapiens α*-synuclein (Uniprot ID P37840), **(B)** *M. truncatula* PM25 (Uniprot ID Q2Q4X9), **(C)** *Human adenovirus 12* Early E1A protein (Uniprot ID P03259), **(D)** *S. cerevisiae* UMP1 (Uniprot ID P38293), and **(E)** *H. sapiens* 4E-BP2 (Uniprot ID Q13542). To the right of each disorder profile is the legend accompanied by a summary of the total percent of predicted disorder for each protein that metapredict V2 (top) or legacy metapredict (bottom) identified. This summary includes the percent of residues identified in each sequence using either the number of amino acids along the sequence with a score greater than or equal to the threshold disorder value (‘Residues over threshold’, left) or the residues identified as disordered using the predict_disorder_domains() function (‘Residues identified in IDR’, right).

Protein residues can be classified as being in IDRs in two different ways. The first involves a binary disorder threshold of 0.5, whereby any residue above a disorder score of 0.5 is considered to be ‘disordered’ and any residue below 0.5 is considered to be ‘ordered’. Using this approach, metapredict V2 did not always identify these proteins as fully disordered; however, it did predict a much larger percentage of the protein to be disordered than legacy metapredict (**Fig. 4C, D, E**, left columns).

Protein residues can also be classified as being inside IDRs using an algorithm to extract contiguous stretches of residues based on a combination of protein heuristics and physical limitations of protein folding (e.g. a folded domain cannot be just a few residues long). This domain identification approach is available via the **`predicted_disorder_domains()`** function, and we have generally found it to be more accurate in identifying discrete boundaries between folded and disordered regions. Indeed, for all five proteins that metapredict legacy failed to identify as disordered, every residue was annotated as being in an IDR with metapredict V2, in line with experimental evidence (**Fig. 4C, D, E**, right columns).

### Improved user experience and interpretability of IDR boundaries

In addition to improved accuracy, metapredict V2 provides a more robust and easy-to-interpret means of predicting contiguous IDRs within proteins. This IDR prediction can be accessed as a command-line tool, as a Python package, and as a web server. In particular, metapredict V2 provides much sharper boundaries between folded and disordered regions, enabling a more robust approach for extracting contiguous disordered and folded domains.

The metapredict web server (https://metapredict.net/) now automatically extracts and colors IDRs, such that IDRs are easily identified from the sequence. In addition, we added a new command-line tool (`metapredict-name`), which enables disorder predictions based on gene names and species. Finally, as a Python package, the function `predict_disorder_domains()` now returns a Python object containing information on the predicted disordered regions and folded domains via dot operator variables. Together, these features all improve the user experience when working with metapredict across the various modalities in which it is implemented.

### Human transcription factors are, on average, over 50% disordered

In the original implementation of metapredict, we noticed that certain types of IDRs yielded false-negative predictions. In particular, transcription factors (TFs) contain IDRs that were often predicted to be partially or entirely folded, despite strong experimental evidence to the contrary. As an example, the transcription factor GCN4 possesses a C-terminal basic zipper (bZIP) DNA binding domain (DBD), but the remainder of the protein is highly disordered in isolation (9, 10). In metapredict legacy, a large swathe of this IDR was predicted to be folded, and the boundary between the DBD and the IDR was incorrectly defined (**Fig. 5A**). In metapredict V2, the entire IDR is correctly identified, and the boundary between this IDR and the DBD is correct to within two residues (**Fig. 5A**).

**Figure 5.**
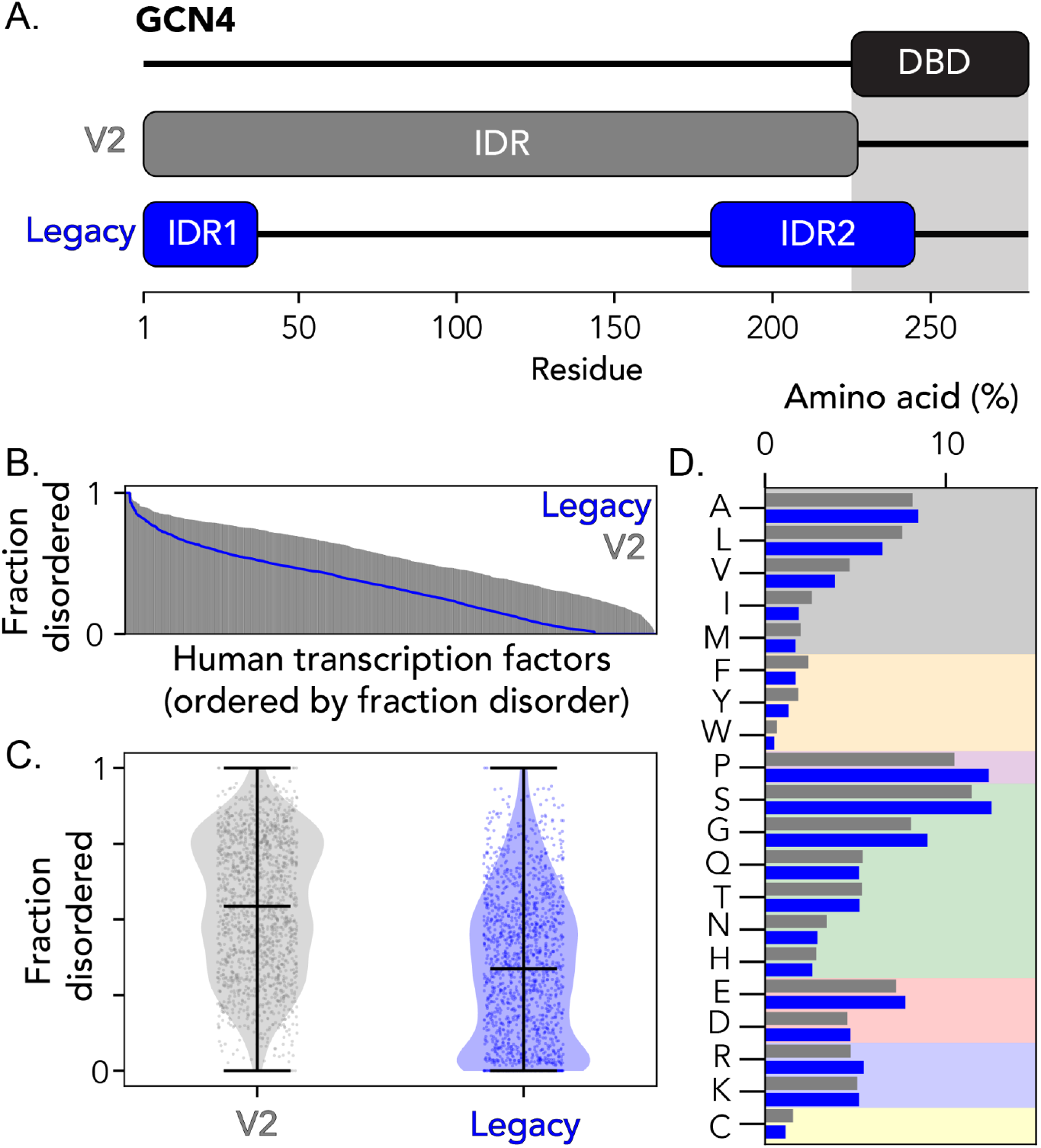
Metapredict V2 provides a more accurate prediction of IDRs in transcription factors. **(A)** Overview of disordered regions in the yeast transcription factor GCN4 (Uniprot ID P03069). Metapredict legacy misses most of the N-terminal portion of the disordered region, while Metapredict V2 correctly identifies this region and the boundary between the folded DNA binding domain (DBD). **(B)** Human transcription factors ordered by the fraction of disordered residues shown for both legacy and V2 versions. **(C)** Fraction of disorder in human transcription factors as calculated using legacy and V2 versions. **(D)** Average amino acid composition for IDRs in human transcription factors as identified by legacy and V2 versions.

Given the improvement in extracting the IDR from GCN4, we sought to determine if the sequence compositions of TF IDRs as predicted by metapredict V2 were distinct from those predicted by metapredict legacy. We used metapredict v2 and legacy metapredict to extract all IDRs from 1,608 human transcription factors (11). Metapredict V2 predicts a much larger fraction of transcription factors to be disordered than legacy metapredict and suggests that on average, transcription factors are over 50% disordered (**Fig. 5B, C**). We interpret this result to reflect the fact that many transcription factors are poised to bind cognate co-activators, leading to sequence features that – in metapredict legacy – made them challenging to distinguish from *bona fide* folded domains (9, 10, 12–14).

An analysis of IDR composition reveals that IDRs identified using metapredict V2 are slightly more hydrophobic than those from metapredict legacy. IDRs from metapredict V2 have a higher fraction of aliphatic and aromatic residues, although we note the differences between legacy and V2 are between 0-2%. Our analysis confirms that human transcription factors are highly disordered in general, while also demonstrating that the composition of transcription factor IDRs does not substantially change when comparing metapredict V2 to legacy metapredict. This suggests that rather than altering the threshold for disorder identification based on composition, metapredict V2 is taking advantage of specific structural patterns encoded through the ppLDDT component of the predictor.

In summary, all available evidence supports the conclusion that metapredict V2 offers enhanced performance and accuracy, especially for sequences that may undergo folding-upon-binding that are typically difficult for canonical predictors to classify correctly.

### Assessment against non-natural proteins

While incorporating ppLDDT information into metapredict has improved accuracy for naturally occurring proteins, a possible concern is that this may have a detrimental effect on non-natural proteins. This could include protein sequences with mutations, *de novo* synthetic proteins, or random peptides synthesized without evolutionary constraints or structure-informed design principles. To explore this possible weakness, we examined examples of each of these three cases.

The polypeptide encoded by exon 1 (Ex1) of the human Huntingtin protein (Uniprot ID P42858) is well-studied in the context of Huntington’s disease (15–18). This ~100-residue protein fragment is an aggregation-prone disordered protein. Ex1 contains a 21-residue glutamine tract that undergoes repeat expansion and this expansion results in an increased propensity for aggregation (18–22). With this in mind, the Q_40_ variant of Ex1 represents a protein with a disease-associated repeat expansion that is not found in the canonical human protein, yet has been well-characterized by a slew of biophysical methods including simulations, circular dichroism spectroscopy, single-molecule spectroscopy, and nuclear magnetic resonance (NMR) spectroscopy (18–25).

Given that coiled-coil domains are often glutamine-rich, we wondered if a Q_40_ variation of Ex1 may mislead metapredict V2 and AlphaFold2 predictions into predicting a folded protein (26). Indeed, structure predictions with AlphaFold2 performed using ColabFold yielded a high-confidence prediction for a single helix extending through the 17-residue N-terminus (which is known to possess transient helicity) into the 40-residue Gln repeat (2, 27) (**Fig. 6A, B**). Encouragingly, however, metapredict V2 correctly predicts this entire sequence to be disordered, with a slight decrease in predicted disorder around the residues known to form a transient helix in the N-terminal fragment (**Fig. 6c**). These results help assure that even for sequences that AlphaFold2 demonstrably and confidently predicted incorrectly, metapredict V2 can delineate disordered regions from folded regions.

**Figure 6.**
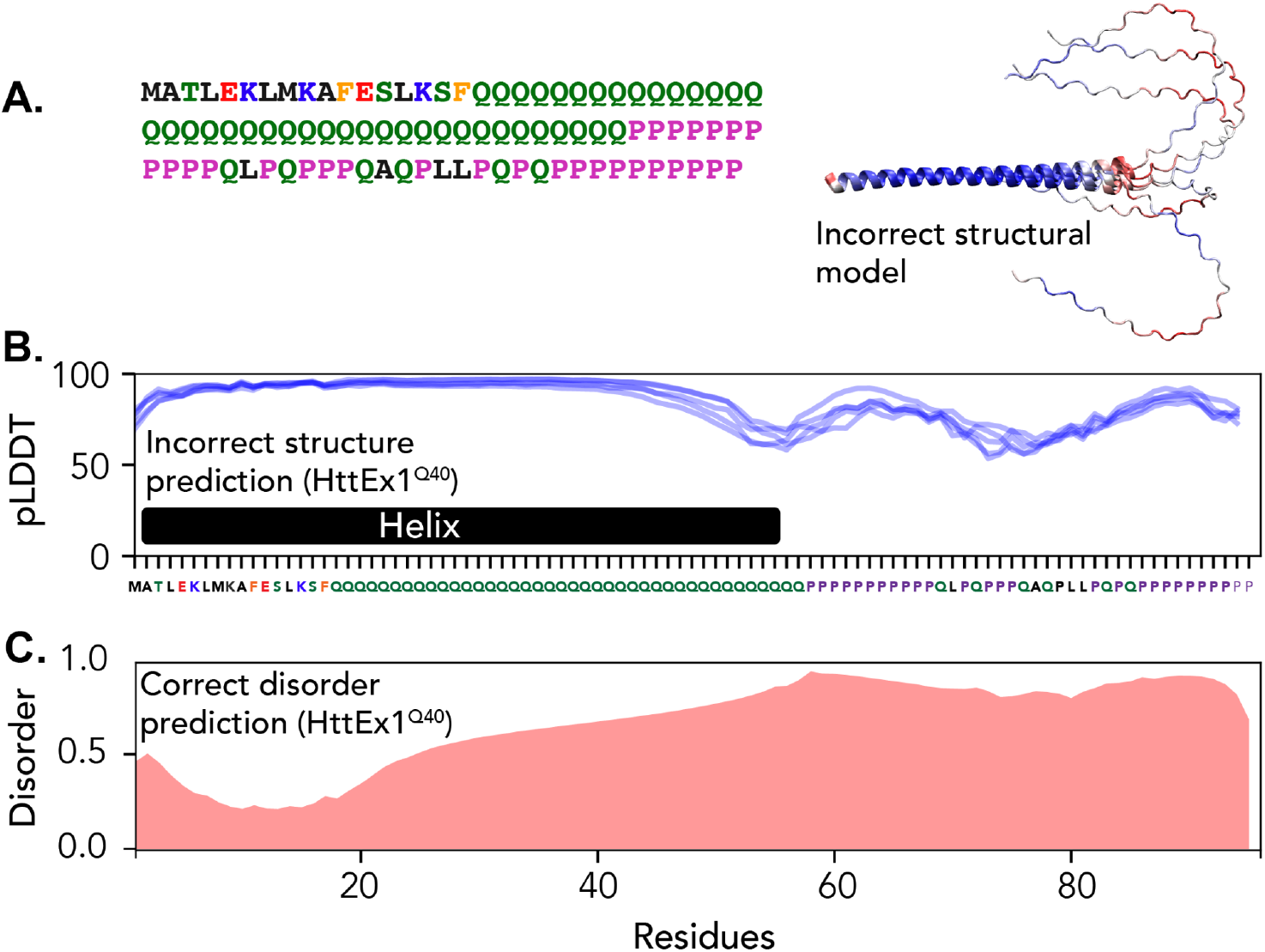
Metapredict V2 correctly predicts a Q_40_ variant of the Huntington Exon1 (Ex1) to be disordered. **(A)** Amino acid sequence of the Q_40_ isoform of Ex1 and predicted structural models from ColabFold-derived AlphaFold2 prediction colored by pLDDT scores (blue is high, red is low). **(B)** Per-residue pLDDT scores from the top five AlphaFold2 models, with the high-confidence helix highlighted. **(C)** Ex1 is predicted to be 100% disordered, although the per-residue disorder dips around the N-terminal region that possesses transient helicity, consistent with ongoing NMR work that suggests this region induces helicity in the polyglutamine tract (23).

We next wondered if metapredict V2 was able to correctly classify *de novo* synthetic proteins. To explore this, we took three protein sequences and structures generated by a deep hallucination network approach for *de novo* sequence design (28). Metapredict V2 is not only able to unambiguously predict all three proteins to be folded but correctly identifies the boundary between the disordered HIS-tag and the N-terminal methionine in the folded domain without error (**Fig. 7**). This analysis suggests that even for *de novo* synthetic proteins, metapredict V2 retains a high level of accuracy.

**Figure 7.**
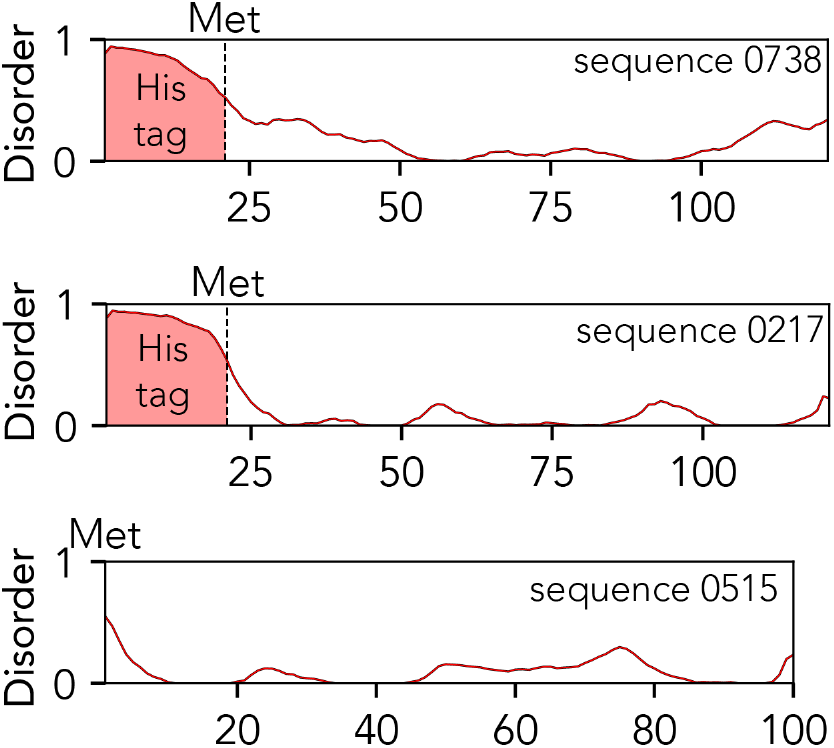
Metapredict V2 correctly predicts the disordered regions in three *de novo* proteins generated by a deep hallucination network (28). In sequences 0738 and 0217, the deposited structure and sequence possess an N-terminal HIS-tag, which metapredict V2 correctly identifies as being disordered, placing the boundary between the HIS-tag and the folded domain at the initiator methionine. In the bottom sequence (sequence 0515) no such HIS tag is present, and no residues are predicted to be disordered.

Finally, we wondered if polypeptides with randomly selected amino acid composition (but no deliberate design rules) could be correctly classified as disordered or ordered with metapredict V2. Tretyachenko *et al.* recently showed that soluble, randomly generated polypeptide sequences can possess secondary structure (29). Specifically, using compositionally-biased starting libraries, the authors generated, expressed, purified, and measured circular dichroism (CD) spectra for 22 random, synthetic proteins. By clustering these sequences into two groups based on the CD spectra, the authors defined two sets of protein sequences that show either disordered or ordered structural tendencies yet are devoid of any evolutionary constraints. With this dataset, we wondered if metapredict would be able to correctly identify the disordered and ordered proteins from the sequence.

We took these 22 sequences, determined the fraction of residues in IDRs, and classified a protein as “disordered” if over 50% of the residues were found in IDRs or “ordered” if not. Encouragingly, metapredict V2 correctly classified all but one of the sequences, while metapredict legacy performed much more poorly (**Fig. 8**). Curiously, the one sequence metapredict V2 appeared to get wrong has some residual helical content (Fig. S2 in (29)) and undergoes a major increase in helical content upon the addition of the helicity-inducing osmolyte 2,2,2-trifluoroethanol (TFE) (29). We speculate that this sequence may be a meta-stable transient helix, poised to undergo a helix-to-coil transition, such that it sits on the boundary between folded and disordered. In summary, metapredict V2 reveals an encouraging ability to delineate between folded and disordered proteins and protein regions, irrespective of the origin of the protein sequence.

**Figure 8.**
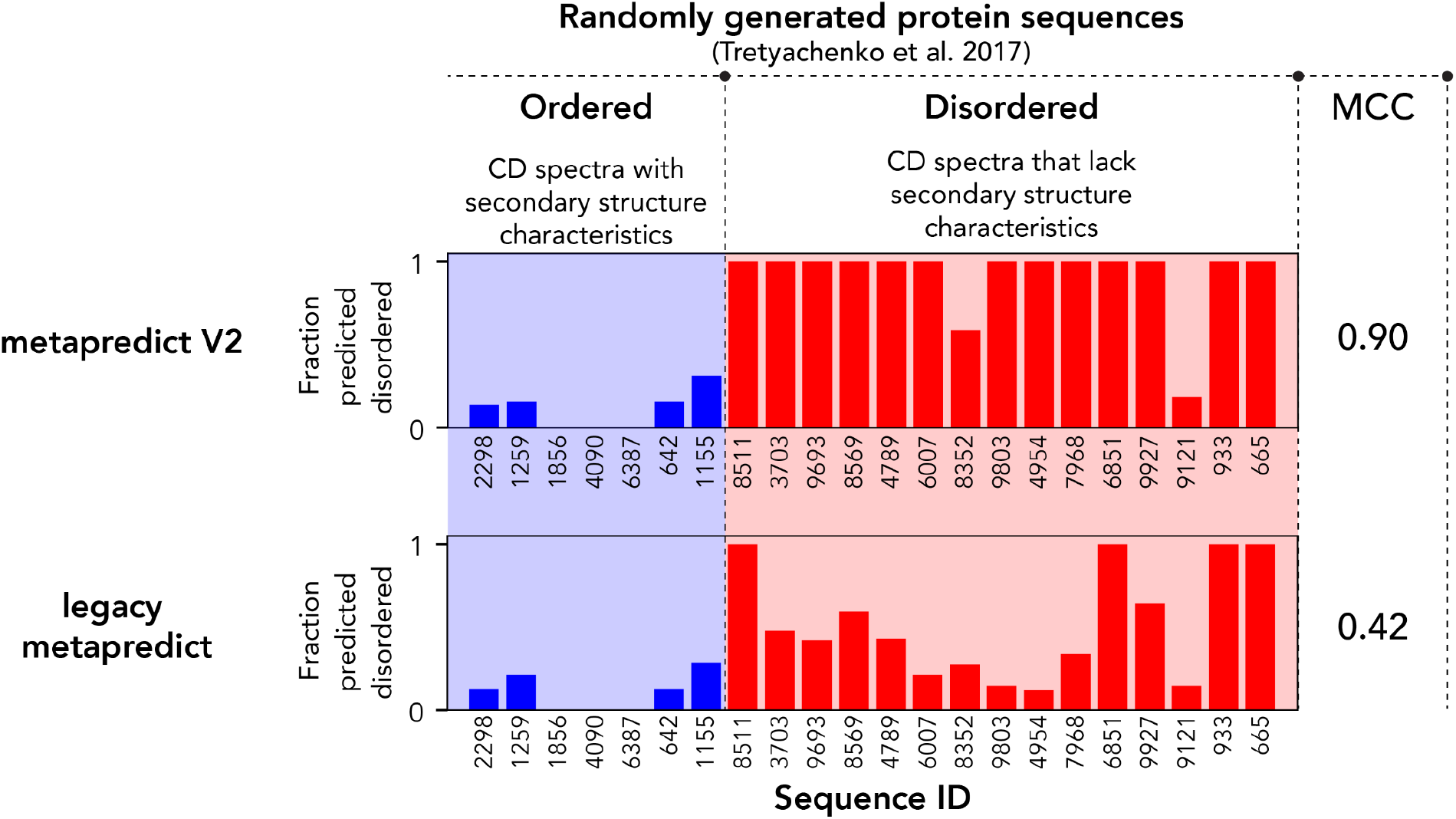
Metapredict V2 correctly classifies randomly generated and experimentally-verified sequences as disordered or ordered. These sequences were taken from (29), and the classification of “ordered” or “disordered’ was based on the clustering of CD spectra with one another. Ordered sequences are those that cluster in a group of CD spectra that possess features consistent with secondary structure elements. The Sequence ID reflects those used in the original paper.

## DISCUSSION

Proteins that are entirely disordered or contain disordered regions play important roles in numerous biological processes (30–32). With this in mind, the ability to quickly, easily, and accurately predict disordered regions from protein sequence information is a central feature in *de novo* sequence analysis. It was this need that led us to develop and implement metapredict, a deep-learning-based meta-predictor of disorder.

Our first implementation of metapredict offered a relatively accurate protein disorder predictor capable of predicting disorders at extremely fast speeds (1). Here we describe an update to metapredict (metapredict V2) that improves prediction accuracy by integrating predicted pLDDT scores (**Fig. 2**, **3**). Although execution times for metapredict V2 are ~50% of that of legacy metapredict, metapredict V2 predicts protein disorder at a rate of between ~6,000 and ~12,000 residues per second, depending on the hardware used (Supplemental Figure 1). To put this in context, metapredict V2 is still capable of predicting the disorder of every protein in the human proteome in 40-60 minutes, depending on the hardware, and remains among the fastest disorder predictors currently available. Importantly, for users who have integrated legacy metapredict into their pipeline or rely on the speed that legacy metapredict provides, this version remains fully accessible both from the command line and from within Python.

Although metapredict V2 is slower than legacy metapredict, we feel that the improved accuracy of metapredict V2 is sufficient to offset this loss in speed. When evaluating the accuracy of metapredict V2 in comparison to 33 other disorder predictors (including legacy metapredict), metapredict V2 was identified as the second most accurate disorder predictor, with the difference between metapredict V2 and the most accurate currently available predictor being just 0.281 residues per 100 predicted (**Fig. 9**). As described previously, metapredict is orders of magnitude faster in execution time compared to the other high-performing predictors (1). As such, we suggest metapredict V2 provides an ideal balance of speed, accuracy, and availability.

**Figure 9.**
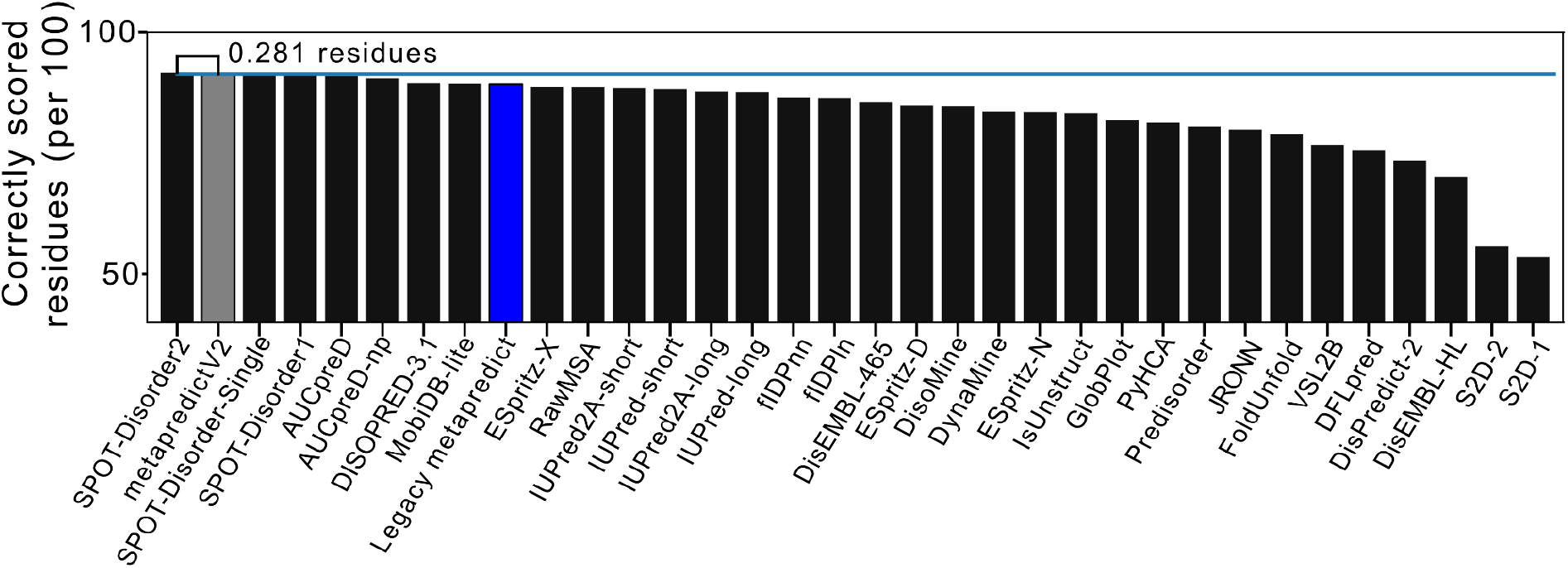
Comparing the accuracy of metapredict V2 and legacy metapredict to other disorder predictors. This chart shows the rank order of predictors in residues correctly scored per 100 residues using true positive and true negative values from the Disprot-PDB dataset. metapredict V2 is highlighted in grey and legacy metapredict is highlighted in blue. To ensure fairness, the per residue scoring here was done using a hard disorder threshold, as opposed to the more `accurate predict_disorder_domains()`.

Despite metapredict V2 showing improved overall accuracy in all tests we have run to date, there are some potential limitations in disorder prediction using this new version of metapredict. These limitations stem primarily from the fact that metapredict V2 works by combining disorder scores from legacy metapredict with predicted pLDDT scores. As mentioned earlier, pLDDT scores are used for AlphaFold2 (AF2) predicted protein structures to quantify how confident one can be in the predicted structure (2). The scores used to make metapredict V2 were *predicted* pLDDT (ppLDDT) scores, which were generated using a BRNN trained on AF2 pLDDT scores from 21 proteomes (1). Thus, if there are any consistent circumstances where AF2 generates a high pLDDT score for a given disordered region or type of disordered region, metapredict V2 will be unlikely to predict the region to be disordered. Indeed, a recent publication highlighted examples where known disordered proteins or protein regions had high AF2 pLDDT scores, which they found to be at least in part due to the disordered regions undergoing conditional folding (4).

Interestingly, when we examined the ability of metapredict V2 to predict the disorder of some proteins identified as having high AF2 pLDDT scores, despite being known to be disordered, we found that metapredict V2 was often able to correctly identify the proteins as being disordered (**Fig. 4A, 4E**). When we examined the predicted pLDDT scores for these proteins, we found that our pLDDT predictor did not give these regions predicted pLDDT scores as high as the actual AF2-generated pLDDT scores. This may suggest that our pLDDT predictor does not always produce high pLDDT scores for some disordered regions, even though the actual AF2 pLDDT scores for the same region are relatively high. However, we note that this is not always the case, as we were able to identify some known disordered protein regions that our pLDDT predictor generated high ppLDDT scores. In particular, AF2 and ppLDDT scores occasionally predict structure for regions predicted to form unreasonably long alpha-helices. Thus, these same proteins were not predicted to be disordered by metapredict V2. Nonetheless, metapredict V2 still offers a substantial improvement in accuracy over legacy metapredict.

## METHODS

### Proteomes used for metapredict V2 network training

21 different proteomes were used to train metapredict V2, as listed below. For each sequence in each proteome, we computed the metapredict hybrid disorder profile. These data were then used as the training/test data for a PARROT-derived BRNN-BRNN network.

UP000002485_284812_SCHPO, UP000000805_243232_METJA, UP000001450_36329_PLAF7, UP000005640_9606_HUMAN, UP000001584_83332_MYCTU, UP000001940_6239_CAEEL, UP000000625_83333_ECOLI, UP000002296_353153_TRYCC, UP000000803_7227_DROME, UP000007305_4577_MAIZE, UP000002195_44689_DICDI, UP000002311_559292_YEAST, UP000002494_10116_RAT, UP000008153_5671_LEIIN, UP000008816_93061_STAA8, UP000006548_3702_ARATH, UP000008827_3847_SOYBN, UP000000437_7955_DANRE, UP000000589_10090_MOUSE, UP000059680_39947_ORYSJ, UP000000559_237561_CANAL

### Metapredict V2 BRNN-LSTM network training

Using PARROT, we trained a bidirectional recurrent neural network with long short-term memory (BRNN-LSTM) on the metapredict hybrid scores. This network was trained as a regression model using one-hot encoding, a hidden vector size of 20, and 2 layers. In addition, we used a learning rate of 0.001, a batch size of 32, and training was done using 200 epochs. The 363,265 sequences derived from the 21 proteomes were split using a 70:15:15 split for training, validation, and testing, respectively.

### Comparing Metapredict V2 with DisProt and CAID

Comparisons between Metapredict V2 and legacy metapredict utilized the same testing from the CAID analysis as was carried out in the original metapredict manuscript (1). Briefly, CAID analyses used the dataset provided by (8). These sequences were obtained from the DisProt database (7). Accuracy was quantified using the Matthew’s Correlation Coefficient (MCC). A cutoff value of 0.5 was used such that a predicted disorder value generated by metapredict V2 with a value greater than or equal to 0.5 was considered disordered whereas a value below 0.5 was not considered to be disordered. The Disprot-PDB dataset was used for accuracy analysis because it contains regions that have been experimentally determined to be either disordered or not disordered, allowing for the identification of true positive, true negative, false positive, and false-negative predictions generated by metapredict V2.

### Sequence analysis

Human transcription factors were taken from Lambert, Jolma, Campitelli *et al. (11)*. Huntingtin protein sequence was taken from https://www.uniprot.org/uniprot/P42858 with polyQ expansion as described in (18). Structural modeling was performed using ColabFold with AlphaFold2 (2, 27, 33). Per residue helicity was computed using the DSSP algorithm as implemented in SOURSOP (https://soursop.readthedocs.io/), a simulation package built on top of MDTraj (34, 35). Sequences and structures for synthetic proteins from deep hallucination design were taken from (28) using PDB IDs 7K3H, 7M0Q, and 7M5T. Sequences for random polypeptide sequences were taken from (29). All sequences and code for sequence analysis figures as well as the full output from the ColabFold prediction of HttEx1^Q40^ are available on our GitHub repository (see below for link).

### Performance tests

All performance tests were carried out using a 2021 14” Apple MacBook Pro with the Apple M1 Max system on a chip (configured with 10 CPU cores and 24 GPU cores) and 64 GB of unified memory. The number of residues per second was calculated by having the predictor (legacy metapredict, metapredict hybrid, or metapredict V2) predict the disorder for 500 randomly generated amino acid sequences of 500 amino acids in length and dividing the total execution time for the prediction to complete by the number of residues predicted.

### Human transcription factor analysis

Human transcription factors were taken from Lambert, Jolma, Campitelli, *et al.* (11). Code, transcription factor FASTA files, and IDRs in SHEPHARD domains format are provided at https://github.com/holehouse-lab/supportingdata/. Specifically, the associated files can be found under the 2021/emenecker_metapredict_2021/v2/transcription_factors/directory. To analyze human transcription factors we recommend using SHEPHARD (https://shephard.readthedocs.io/en/latest/overview.html).

## Code availability

All code used in the original metapredict paper has been updated to work with metapredict V2 and can be found at https://github.com/holehouse-lab/supportingdata/under the 2021/emenecker_metapredict_2021/directory.

## Acknowledgments

This work was supported by NSF IntBio-MCB-2128068 to A.S.H. and NSF Graduate Research Fellowship DGE-1247271 and DGE-2139839 to D.G. We are extremely grateful for helpful comments and suggestions from Aidan Flynn. We thank various members of the IDP community for fruitful discussions as we have continued to enhance and improve metapredict. A.S.H. is a scientific consultant with Dewpoint Therapeutics. All other authors declare no competing interests.

## Notes

https://github.com/holehouse-lab/supportingdata/tree/master/2021/emenecker_metapredict_2021/v2

### Competing Interest Statement

A.S.H. is a scientific consultant for Dewpoint Therapeutics

### Summary of Updates

Fixed some typos, improved the text, all cosmetic, no scientific changes. Thanks Aidan!

